## Supplemental Figures for "DNA Methylation-Based High-Resolution Mapping of Long-Distance Chromosomal Interactions in Nucleosome-Depleted Regions"

### Slide 1
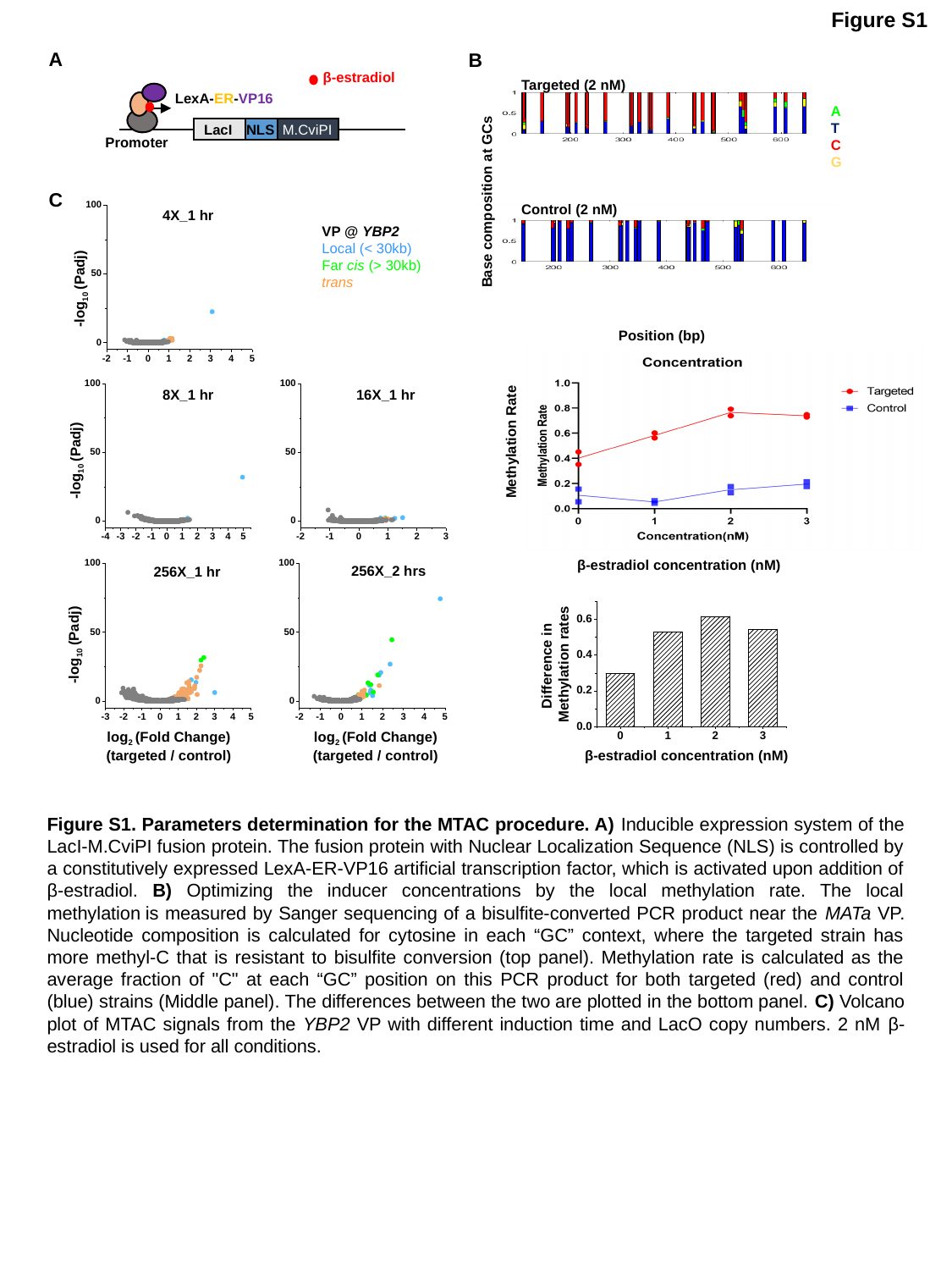

Figure S1
A
B
β-estradiol
LexA-ER-VP16
LacI
NLS
M.CviPI
Promoter
Targeted (2 nM)
Control (2 nM)
Base composition at GCs
Position (bp)
A
T
C
G
C
4X_1 hr
VP @ YBP2
Local (< 30kb)
Far cis (> 30kb)
trans
-log_10_ (Padj)
8X_1 hr
16X_1 hr
-log_10_ (Padj)
256X_2 hrs
256X_1 hr
-log_10_ (Padj)
log_2_ (Fold Change)
(targeted / control)
log_2_ (Fold Change)
(targeted / control)
 Methylation Rate
β-estradiol concentration (nM)
Difference in  Methylation rates
β-estradiol concentration (nM)
Figure S1. Parameters determination for the MTAC procedure. A) Inducible expression system of the LacI-M.CviPI fusion protein. The fusion protein with Nuclear Localization Sequence (NLS) is controlled by a constitutively expressed LexA-ER-VP16 artificial transcription factor, which is activated upon addition of β-estradiol. B) Optimizing the inducer concentrations by the local methylation rate. The local methylation is measured by Sanger sequencing of a bisulfite-converted PCR product near the MATa VP. Nucleotide composition is calculated for cytosine in each “GC” context, where the targeted strain has more methyl-C that is resistant to bisulfite conversion (top panel). Methylation rate is calculated as the average fraction of "C" at each “GC” position on this PCR product for both targeted (red) and control (blue) strains (Middle panel). The differences between the two are plotted in the bottom panel. C) Volcano plot of MTAC signals from the YBP2 VP with different induction time and LacO copy numbers. 2 nM β-estradiol is used for all conditions.

### Slide 2
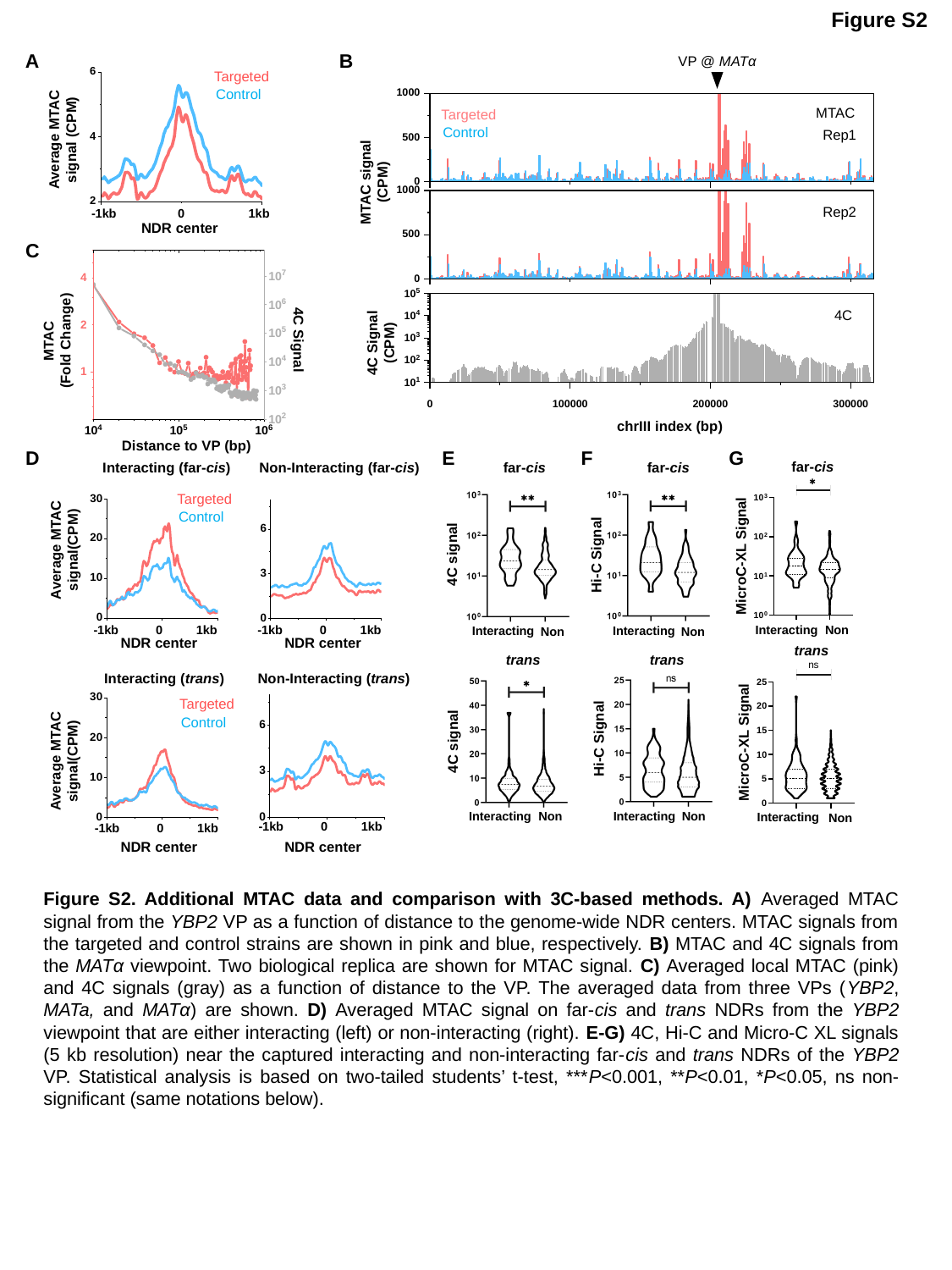

Figure S2
A
B
VP @ MATα
MTAC
Targeted
Control
Rep1
MTAC signal (CPM)
Rep2
4C
4C Signal (CPM)
chrIII index (bp)
Average MTAC signal (CPM)
0
-1kb
1kb
NDR center
Targeted
Control
C
4
2
MTAC
(Fold Change)
4C Signal
1
Distance to VP (bp)
G
D
E
F
far-cis
MicroC-XL Signal
Interacting
Non
far-cis
far-cis
4C signal
Hi-C Signal
Interacting
Interacting
Non
Non
trans
MicroC-XL Signal
Interacting
Non
trans
trans
Hi-C Signal
4C signal
Interacting
Non
Interacting
Non
Interacting (far-cis)
Non-Interacting (far-cis)
Targeted
Control
Average MTAC signal(CPM)
-1kb 0 1kb
-1kb 0 1kb
NDR center
NDR center
Non-Interacting (trans)
Interacting (trans)
Targeted
Control
Average MTAC signal(CPM)
-1kb 0 1kb
-1kb 0 1kb
NDR center
NDR center
Figure S2. Additional MTAC data and comparison with 3C-based methods. A) Averaged MTAC signal from the YBP2 VP as a function of distance to the genome-wide NDR centers. MTAC signals from the targeted and control strains are shown in pink and blue, respectively. B) MTAC and 4C signals from the MATα viewpoint. Two biological replica are shown for MTAC signal. C) Averaged local MTAC (pink) and 4C signals (gray) as a function of distance to the VP. The averaged data from three VPs (YBP2, MATa, and MATα) are shown. D) Averaged MTAC signal on far-cis and trans NDRs from the YBP2 viewpoint that are either interacting (left) or non-interacting (right). E-G) 4C, Hi-C and Micro-C XL signals (5 kb resolution) near the captured interacting and non-interacting far-cis and trans NDRs of the YBP2 VP. Statistical analysis is based on two-tailed students’ t-test, ***P<0.001, **P<0.01, *P<0.05, ns non-significant (same notations below).

### Slide 3
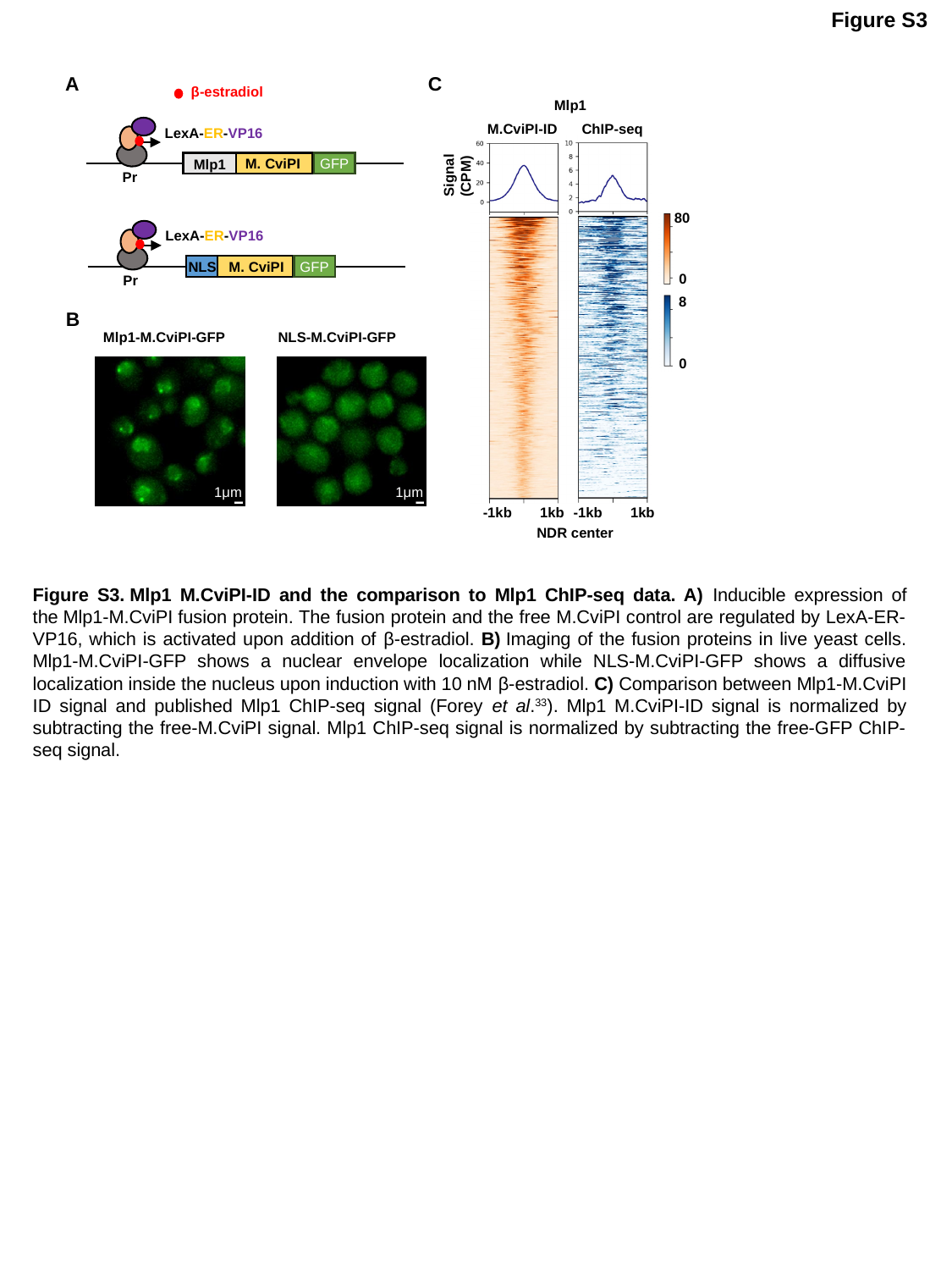

Figure S3
C
A
β-estradiol
LexA-ER-VP16
M. CviPI
Mlp1
Pr
Mlp1
ChIP-seq
M.CviPI-ID
Signal (CPM)
80
0
8
0
-1kb 1kb
-1kb 1kb
NDR center
GFP
LexA-ER-VP16
NLS
M. CviPI
GFP
Pr
B
Mlp1-M.CviPI-GFP
NLS-M.CviPI-GFP
1μm
1μm
Figure S3. Mlp1 M.CviPI-ID and the comparison to Mlp1 ChIP-seq data. A) Inducible expression of the Mlp1-M.CviPI fusion protein. The fusion protein and the free M.CviPI control are regulated by LexA-ER-VP16, which is activated upon addition of β-estradiol. B) Imaging of the fusion proteins in live yeast cells. Mlp1-M.CviPI-GFP shows a nuclear envelope localization while NLS-M.CviPI-GFP shows a diffusive localization inside the nucleus upon induction with 10 nM β-estradiol. C) Comparison between Mlp1-M.CviPI ID signal and published Mlp1 ChIP-seq signal (Forey et al.^33^). Mlp1 M.CviPI-ID signal is normalized by subtracting the free-M.CviPI signal. Mlp1 ChIP-seq signal is normalized by subtracting the free-GFP ChIP-seq signal.

### Slide 4
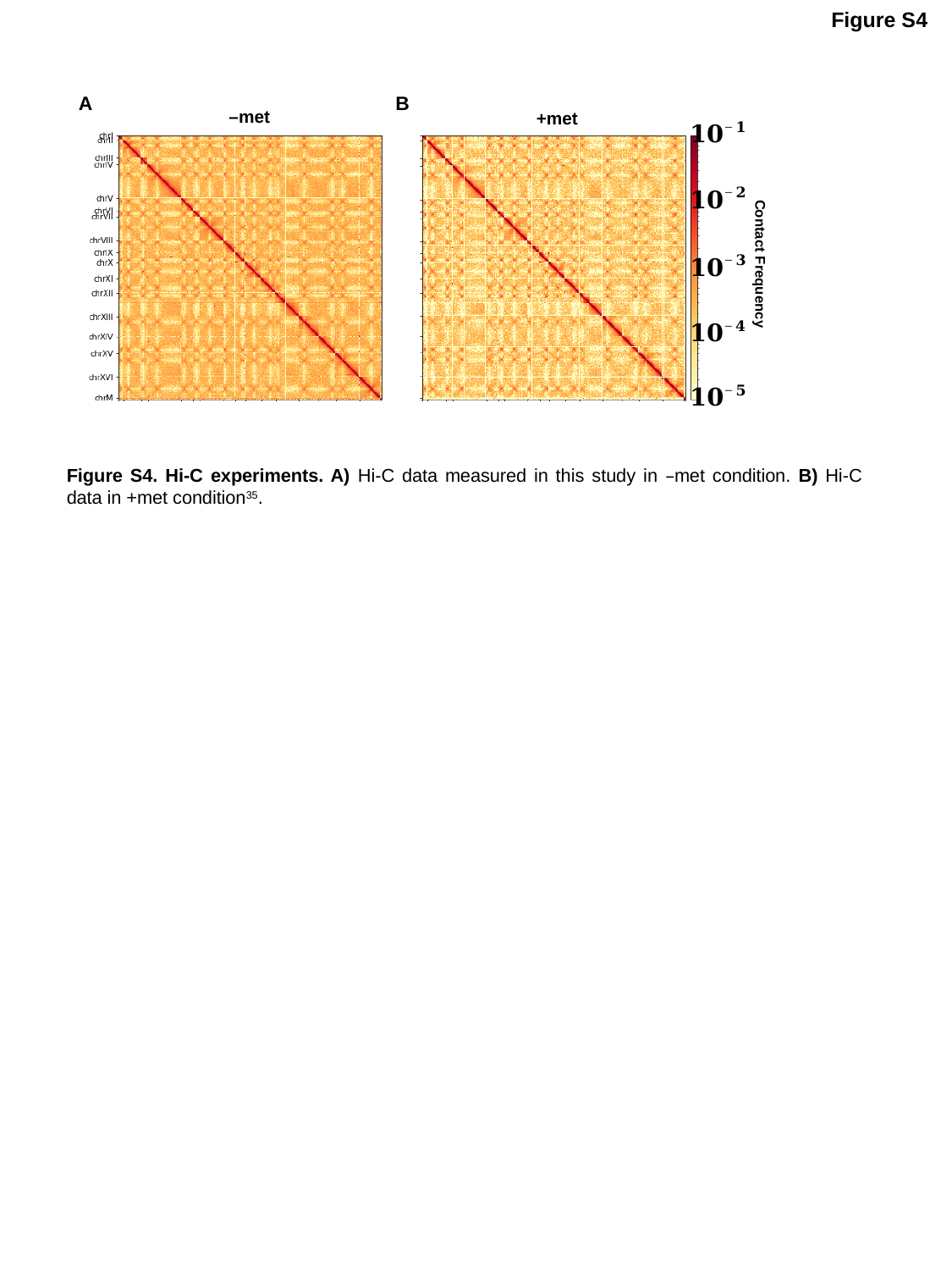

Figure S4
A
B
–met
+met
Contact Frequency
Figure S4. Hi-C experiments. A) Hi-C data measured in this study in –met condition. B) Hi-C data in +met condition^35^.

### Slide 5
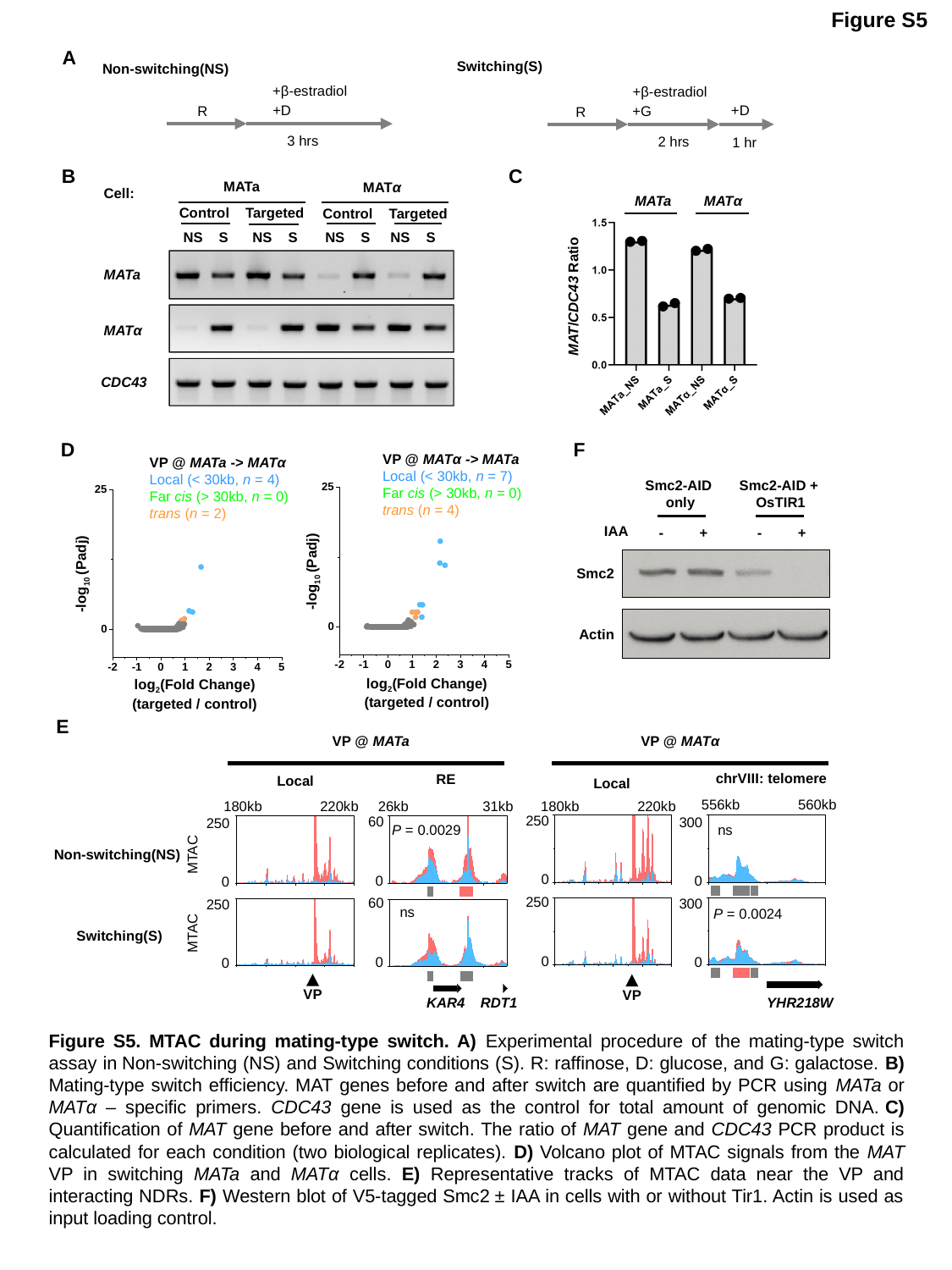

Figure S5
A
Switching(S)
Non-switching(NS)
+β-estradiol
+D
R
3 hrs
+β-estradiol
+D
+G
2 hrs
1 hr
R
B
C
MATa
MATα
Cell:
MATa
MATα
MAT/CDC43 Ratio
Control Targeted
Control Targeted
NS S NS S
NS S NS S
MATa
MATα
CDC43
D
F
VP @ MATα -> MATa
Local (< 30kb, n = 7)
Far cis (> 30kb, n = 0)
trans (n = 4)
-log_10_ (Padj)
log_2_(Fold Change)
(targeted / control)
VP @ MATa -> MATα
Local (< 30kb, n = 4)
Far cis (> 30kb, n = 0)
trans (n = 2)
-log_10_ (Padj)
log_2_(Fold Change)
(targeted / control)
Smc2-AID
only
Smc2-AID +
OsTIR1
IAA
- +
- +
Smc2
Actin
E
VP @ MATα
VP @ MATa
chrVIII: telomere
RE
Local
Local
560kb
556kb
31kb
26kb
220kb
220kb
180kb
180kb
250
 60
300
250
ns
P = 0.0029
Non-switching(NS)
MTAC
0
0
0
0
250
 60
300
250
ns
P = 0.0024
MTAC
Switching(S)
0
0
0
0
VP
VP
RDT1
YHR218W
KAR4
Figure S5. MTAC during mating-type switch. A) Experimental procedure of the mating-type switch assay in Non-switching (NS) and Switching conditions (S). R: raffinose, D: glucose, and G: galactose. B) Mating-type switch efficiency. MAT genes before and after switch are quantified by PCR using MATa or MATα – specific primers. CDC43 gene is used as the control for total amount of genomic DNA. C) Quantification of MAT gene before and after switch. The ratio of MAT gene and CDC43 PCR product is calculated for each condition (two biological replicates). D) Volcano plot of MTAC signals from the MAT VP in switching MATa and MATα cells. E) Representative tracks of MTAC data near the VP and interacting NDRs. F) Western blot of V5-tagged Smc2 ± IAA in cells with or without Tir1. Actin is used as input loading control.

### Slide 6
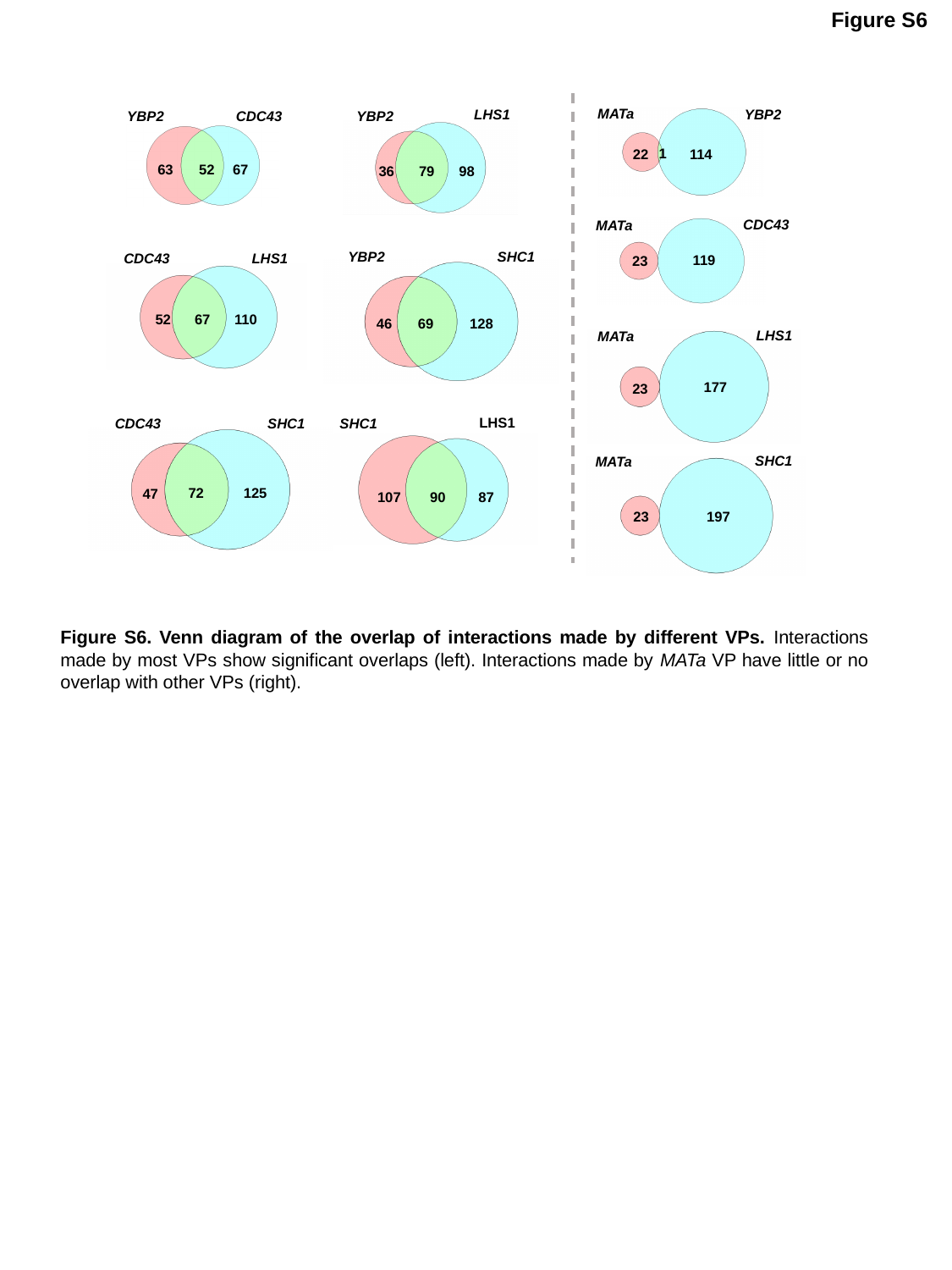

Figure S6
MATa
YBP2
1
114
22
LHS1
YBP2
36
79
98
YBP2
CDC43
52
67
63
CDC43
MATa
119
23
SHC1
YBP2
69
128
46
CDC43
LHS1
67
110
52
LHS1
MATa
177
23
LHS1
SHC1
90
87
107
SHC1
CDC43
72
125
47
SHC1
MATa
197
23
Figure S6. Venn diagram of the overlap of interactions made by different VPs. Interactions made by most VPs show significant overlaps (left). Interactions made by MATa VP have little or no overlap with other VPs (right).
